# Higher-order thalamic nuclei facilitate the generalization and maintenance of spike-and-wave discharges of absence seizures

**DOI:** 10.1101/2022.12.02.518853

**Authors:** Zoe Atherton, Olivér Nagy, Lívia Barcsai, Péter Sere, Nikolett Zsigri, Tamás Földi, Antal Berényi, Vincenzo Crunelli, Magor L. Lőrincz

**Affiliations:** Neuroscience Division, School of Bioscience, Cardiff University, Cardiff, UK; Department of Physiology, University of Szeged, Szeged 6720, Hungary MTA-SZTE; ‘Momentum’ Oscillatory Neuronal Networks Research Group, Department of Physiology, University of Szeged, Szeged 6720, Hungary; Center for Neural Science, New York University, New York, NY 10016, USA; HCEMM-SZTE Magnetotherapeutics Research Group, University of Szeged; Szeged, 6720, Hungary; Department of Physiology, Anatomy and Neuroscience, Faculty of Sciences, University of Szeged, Szeged, Hungary

**Keywords:** Epilepsy, somatosensory cortex, thalamus, higher-order thalamic nuclei, ensemble recordings

## Abstract

Spike-and-wave discharges (SWDs), generated by the cortico-thalamo-cortical (CTC) network, are pathological, large amplitude oscillations and the hallmark of absence seizures (ASs). SWDs begin in a cortical initiation network in both humans and animal models, including the Genetic Absence Epilepsy Rats from Strasbourg (GAERS), where it is located in the primary somatosensory cortex (S1). The behavioral manifestation of an AS occurs when SWDs spread from the cortical initiation site to the whole brain, however, the mechanisms behind this rapid propagation remain unclear. Here we investigated these processes beyond the principal CTC network, in higher-order (HO) thalamic nuclei (lateral posterior (LP) and posterior (PO) nuclei) since their diffuse connectivity and known facilitation of intracortical communications make these nuclei key candidates to support SWD generation and maintenance. In freely moving GAERS, multi-site LFP in LP, PO and multiple cortical regions revealed a novel feature of SWDs: during SWDs there are short periods (named SWD-breaks) when cortical regions far from S1, such the primary visual cortex (V1), become transiently unsynchronized from the ongoing EEG rhythm. Inactivation of HO nuclei with local muscimol injections or optogenetic perturbation of HO nuclei activity increased the occurrence of SWD-breaks and the former intervention also increased the SWD propagation-time from S1. The neural underpinnings of these findings were explored further by silicon probe recordings from single units of PO which uncovered two previously unknown groups of excitatory neurons based on their burst firing dynamics at SWD onset. Moreover, a switch from tonic to burst firing at SWD onset was shown to be an important feature since it was much less prominent for non-generalized events, i.e. SWDs that remained local to S1. Additionally, one group of neurons showed a reverse of this switch during SWD-breaks, demonstrating the importance of this firing pattern throughout the SWD. In summary, these results support the view that multiple HO thalamic nuclei are utilized at SWD onset and contribute to cortical synchrony throughout the paroxysmal discharge.

## INTRODUCTION

Absence seizures (ASs) are characterized by a transient lapse in consciousness and are present in several epilepsies including childhood absence epilepsy (CAE) (Crunelli et al., 2020; Panayiotopoulos et al., 1989). Despite a seizure remission rate of 60% in CAE, the impact of ASs remains evident in other facets of life as this epilepsy is co-morbid with social and learning difficulties as well as neuropsychiatric disorders (Caplan et al., 2008) including anxiety, depression and learning deficits (Caplan et al., 2005; Vega et al., 2011). Moreover, approximately 30% of CAE patients are refractory to monotherapy with current first-choice treatments including ethosuximide, valproate and lamotrigine (Glauser et al., 2013), each with unwanted side-effects. The need for more effective treatments is clear and improving our understanding of the epileptogenic and ictogenic mechanisms of ASs will make this goal more achievable.

Spike-and-wave discharges (SWDs), the EEG hallmark of ASs, are thalamo-cortical driven oscillations that originate in cortical initiation sites (Bai et al., 2010; Benuzzi et al., 2012; Tenney et al., 2013; Westmijse et al., 2009) before propagating rapidly across the brain. This phenomenon was originally observed in multiple rat models, where the initiation site is the primary somatosensory cortex (S1) (Meeren et al., 2002; Polack et al., 2007). Due to the location of SWD initiation, the majority of research in animal models has been conducted on the somatosensory thalamo-cortical module including S1, the first-order (FO) ventrobasal thalamus (VB) and the inhibitory nucleus reticularis thalami (NRT). There are numerous changes in this network at SWD onset including increased cortico-cortical associations, led by S1 (Meeren et al., 2002) and feedforward inhibition to the FO VB nucleus via the NRT (McCafferty et al., 2018).

However, this thalamic-cortical module also includes the higher-order (HO) thalamic nucleus, the posterior thalamic group (PO), which has recently come into focus. Being a HO nucleus, the PO receives direct driving input from deep layers of S1 (Sherman and Guillery, 1998) and even prior to SWD onset, there is an increased bidirectional communication between these regions (Lüttjohann and Pape, 2019) that is maintained for the first 500 ms of a seizure (Lüttjohann and van Luijtelaar, 2012). This feature has not been observed in the VB or NRT and thus could be unique to PO or reflect a potential reverberant excitatory role of HO nuclei in general (Lüttjohann and Pape, 2019; Lüttjohann and van Luijtelaar, 2012). Additionally, HO projections are not specific like FO nuclei (Sherman and Guillery, 1998) since PO neurons project not only back to S1, but also to the primary motor cortex (M1) and higher-cortical areas (Casas-Torremocha et al., 2019; Ohno et al., 2012). These features make HO nuclei serious candidates to facilitate synchronization across the cortex but the extent to which they do this remains unknown.

Using freely-moving Genetic Absence Epilepsy Rats from Strasbourg (GAERS), a well-established polygenic model of ASs (Depaulis et al., 2016; Vergnes et al., 1986), this study provides causal evidence, via local pharmacological inhibition and selective optogenetic manipulation, that LP and PO facilitates the propagation and maintenance of SWDs from S1 to M1 and the primary visual cortex (V1). Moreover, single-unit recordings revealed that PO neurons are heterogeneous in their dynamics throughout SWDs and that the burst firing pattern is closely related to SWD onset and is utilized during SWD maintenance. Overall, this work highlights that multiple HO nuclei are potential targets to prevent the generalization and thus, the behavioral manifestation of ASs.

## METHODS

### Animals

Experiments on male GAERS were undertaken in two establishments, the School of Biosciences, Neuroscience Division, Cardiff University, UK and the Department of Physiology, University of Szeged, Hungary. In both institutions, the GAERS were bred in-house with access to food and water *ad libitum*. Both locations had a 12:12 hour light:dark cycle and controlled temperature (19-21°C) and humidity (45-65%). All experiments were conducted on rats aged 4-8 months (under general guidance from recent guidelines on epilepsy animal experimentation) (Lidster et al., 2016) and with approval of the local ethical committees and under the regulations of either the UK Home Office or EU directives.

### Surgery and electrodes

All surgical procedures took place under isoflurane anaesthesia, induced and maintained at 5% and 2% in 2L oxygen/minute, respectively. For cortical LFP recordings, tungsten electrode doublets were implanted at the following coordinates (all mm from Bregma): mPFC (AP: +3.7, ML: 1.3, DV: 3.2), M1 (AP: +3.0, ML: 3.0, DV: 0.8-1.4), V1 (AP: -7.5, ML: 3.0, DV: 0.8-1.4), S1 (AP: +0.2, ML: 5.5, DV: 1.1-1.8), A1 (AP: -5.5, ML: 4.2, DV: 4.5). A HO thalamic electrode targeted AP: -3.6, ML: 2.4, DV: -3 and -4mm to record LP and PO.

For muscimol experiments, micro-infusion cannulae (Bilaney, UK) were implanted bilaterally at AP: -3.6, ML: ± 3.2, DV: 3mm with an angle of 15°. To record thalamic single units, a 4-shank H-32 silicon probe (NeuroNexus, Texas, US) was implanted above PO (AP: -3.6, ML: 2.4, DV: 3.6) and lowered into the target area via a hand-made microdrive adapted from Vandecasteele et al (2012). For optogenetic experiments, rats were bilaterally injected with 500 nl of AAV5-CAMK2-ChR2-TdTom in PO (AP: -3.5, ML: 2.5, DV: 5.2-4.9) and optical fibers (0.2 mm diameter, 0,39 NA, (FT200EMT, Thorlabs) implanted above the injection sites (AP: -3.5, ML: 2.5, DV: 4.9). Gold screws were used for ground and reference electrodes over the cerebellum.

### Data Acquisition

Animals were placed in plexiglass boxes or recorded directly from their home cages for the LFP and single unit recordings, respectively. For the LFP and microinjection experiments, signals were acquired via amplifiers (SuperTech, HU) with a 0.08 - 500Hz band pass filter. Data were digitised at 1 kHz by a 1401 interface (CED, UK) and acquired with Spike2 software (CED, UK). The signals in the single unit and optogenetic experiment were acquired via HS3 headstages (Amplipex, HU), which digitised the signal at 20kHz and amplified it by 200X. The signal then passed to a KJE-1001 amplifier (Amplipex, HU) and acquired via AmpliRec (Amplipex, HU).

### Microinfusions

500 nL of artificial cerebrospinal fluid (aCSF) (Bilaney, UK), which contained the following (in mM): Na^+^ 150; K^+^ 3.0; Ca^2+^ 1.4; Mg^2+^ 0.8; P 1.0; Cl^-^ 155), was injected with or without 15 ng (0.0002 mg/ml) of muscimol (Tocris, UK) into LP or PO via internal cannula (Bilaney, UK) that extended 1 or 2mm beyond the guide cannula, respectively. The experiment was counter-balanced so that half the animals received injection into LP and, after a 3-day wash-out period, had an injection into PO, or vice versa.

### Optogenetic stimulation

Three weeks after vector injection and implantation, the LFP was recorded in the supragrannular layers at the following cortical areas: mPFC, M1, S1, A1 and V1. PO photostimulation consisted of a single light flash (duration: 10 ms, light intensity at the fiber tip: 1-5.5 mW) delivered 5 seconds after a SWD detected in S1. Trials with spontaneous V1 gaps were excluded from the analysis.

### Histological processing

Electrolytic lesions were performed on the last day of the recording experiments to visualise LFP electrodes and silicon probe shanks via a DS3 Isolated Current Stimulator (Digitimer, USA) emitting a total 20µA (at a rate of 1µA/s). The probe tract was stained by application of DiI labelling solution (Thermofisher, USA) before implantation. The location of cannulae was confirmed by injecting a DiI after the final recording and muscimol with a fluorophore conjugate (Thermofisher, US) was used in some animals to test the spread of the drug. Animals were sacrificed with pentobarbital (200mg/kg, Vetoquinol, FR) 24-48 hours later and the brains were stored in 4% PFA for at least 24 hours before slicing at 80mM and imaged with a confocal microscope.

### Seizure extraction

SWDs were extracted via MATLAB: first SWC times were determined based on amplitude threshold crossings and then a SWD was classified as a series of SWCs occurring at the right frequency (i.e. 6-8 SWCs in one second) and duration (i.e. at least 1 second). Since non-generalized events were often short and of a smaller amplitude, they were extracted manually.

### LFP analysis

To explore correlations in amplitude, ictal signals were squared and then an amplitude envelope of the squared signal underwent cross-correlation analysis. The *xcovariance* function was used which normalizes and removes the mean from the signal before the cross-correlation. Lag times were compared with ANOVA between HO and V1 and a t-test was used to test lag differences from zero.

All statistics were completed using R software. Regional differences in each seizure parameter, such as onset time, was completed using a one-way ANOVA.

### Microinfusion analysis

The test period after drug injection (120 min) was split into 30-minute intervals. These intervals were all statistically compared to seizure activity during the 60-minute pre-injection period. The parameters affected by time bin duration, i.e. number of seizures, time spent in seizures and number of non-generalized events, where normalized to the bin duration (i.e. halving values collected in pre-drug period).

### Seizure frequency and power

Power was analyzed by extracting a finite sample of 2 s at the beginning of the seizure, based on SWD times of V1 to ensure all channels were expressing SWCs during this epoch. Any sample that included a seizure break in either M1 or V1 was removed. The percentage change from the baseline was then calculated to generate a mean percentage change at each interval.

### Statistics of microinfusions

The main interest in the microinjection experiment was the effect of muscimol vs aCSF in the two regions, LP and PO. No differences were observed between aCSF injection in LP (n=2) and PO (n=5) and therefore these data were combined to form the control dataset.

For intra-seizure parameters (duration, delay duration, offset difference, SWC-Failure, break rate and break duration and power), the lmer function, from the lme4 package (Bates et al., 2015), was used to produce a linear mixed effect model with the formula:

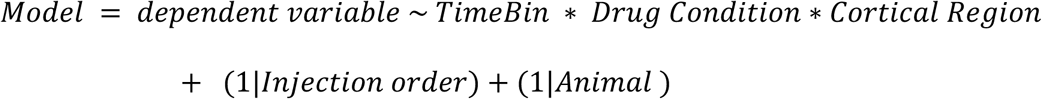

where TimeBin and Drug Condition (i.e. aCSF injection vs muscimol in LP vs. muscimol in PO) were the fixed factors of interest. This allowed many effects to be assessed, across and within condition groups, within a single model. Multiple comparisons were made using lsmeans with a Bonferroni correction.

### Optogenetic analysis

Changes in LFP rhythmicity was quantified by comparing autocorrelations from two epochs: a 1 second pre-stimulation control period and a 1 s epoch post-stimulation. A correlation index was calculated by the side to middle peak ratio and pre- and post-stimulation differences were quantified using a Wilcoxon-signed rank test.

### Single unit analysis

Kilosort 2 (Pachitariu et al., 2016) was used for spike sorting and the clusters were manually refined using the phy package (Cyrille Rossant, 2019). Neuronal firing was classified into total, tonic and burst firing, based on definitions used in McCafferty et al. (2018), and burst profiles were generated to confirm a decelerando and accelerando-decelerando pattern, for TC and NRT neurons, respectively. The firing rate was calculated in 500 ms epochs that slides by 10 ms. The average firing rate was determined for each neuron individually and then an average across neurons was calculated for the population average. Units were classified into groups based on their burst firing dynamics using a guassian mixed model with a seed of 2 groups, based on observation.

### SWC-spike triggered unit spike average

The relation of unit firing to the ongoing SWC-spikes in S1 and V1 was assessed. SWC-spikes were extracted in Spike2 software (Cambridge Electronic Design, UK) by extracting peaks over an amplitude threshold with minimum step of 0.07 s. Spike time difference to each SWC-spike was extracted. Only the first spike (i.e. the spike closest to zero) was considered to understand when PO neurons were first or last activated following or preceding the SWC-spike, respectively. However, since this approach is affected by coherence and amount of neuronal firing, a cumulative distribution function of the same data was also completed.

### Firing rate changes and statistics

The total tonic and burst firing rates were aligned to a time point of interest, i.e. SWD onset or offset, V1 SWD-break onset or onset of S1-only and S1+M1-only SWD. Firing rates were plotted using MATLAB and the *shadedErrorBar* function (Rob Campell, Github: raacampbell). The rate at particular time-points was extracted (detailed in relevant results section) for statistical analysis. Unless stated otherwise, a repeated measures ANOVA was completed on each firing type separately using *anova* function in R. A pairwise post-hoc comparison Tukey test with Bonferronni correction was computed to explore differences within groups, across time, and between unit groups at different time points. This was completed after the assumptions of normality and homoscedasticity were confirmed. If the assumptions were violated, which was the case in some analysis of burst firing, the data was transformed to its log before statistical analysis.

### Autocorrelations

Spikes from all V1-SWD breaks within a neuron were concatenated and a normalised autocorrelogram was obtained via the xcorr function and then averaged across all neurons. The maximum coeeffienct values were obtained for the first and second non-zero peak for each neuron at each epoch and peak differences were obtained relative to value at SWD-break and then averaged. A t-test was used to determine whether peak differences were different to 0.

## RESULTS

### Propagation time of SWDs was longer for cortical regions furthest from S1

LFPs from multiple cortical (S1, M1 and V1) and HO thalamic regions (LP and PO) were recorded in freely-moving GAERS to assess generalization features of SWDs. In 174 SWDs from 6 GAERS (29 ± 5), we observed a duration of 25.3 ± 3.1s and a frequency of 7.2 ± 0.3Hz, alike those previously observed (Depaulis et al., 2016; McCafferty et al., 2018). There was a delay between the appearance of SWDs in S1 and other regions (Figure 1A), hereafter called onset delay, confirming the presence of an initiation site in our sample. Onset delays from S1 to V1 were significantly longer than to all other brain areas (V1: 0.8 ± 0.09 vs. M1: 0.4 ± 0.05sec, p<0.0001; vs LP: 0.4 ± 0.09sec, p<0.0001; vs PO:0.4 ± 0.05 p<0.0001) (Figure 1C).

**Figure 1.**
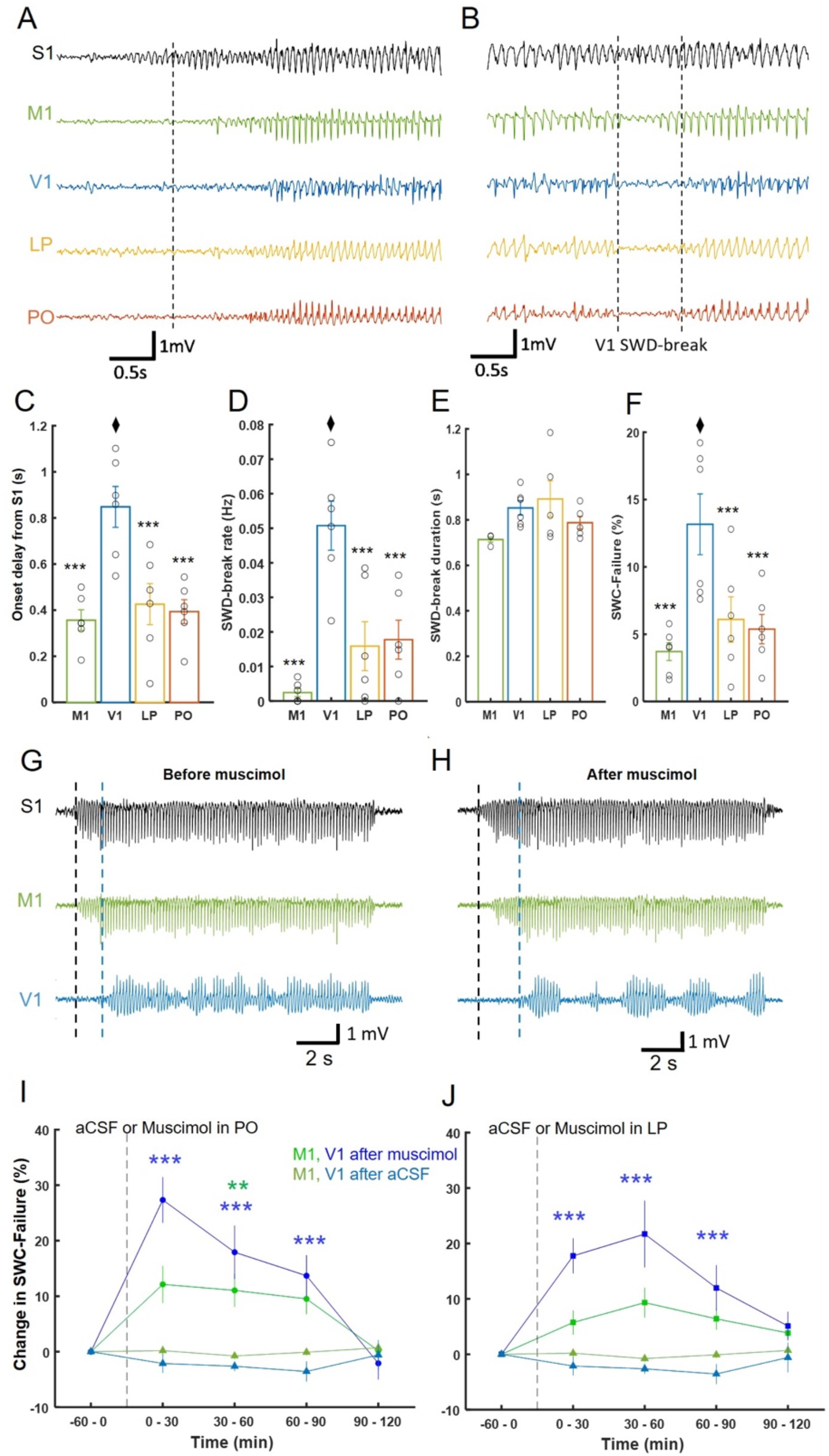
SWD onset delays and SWD breaks contribute to SWC-Failure: effect of pharmacological inhibition of LP and PO. A) LFP traces from cortical regions S1, M1 and V1, and HO thalamic nuclei LP and PO show an onset delay from S1 to the other brain regions. Vertical dashed line indicates the start of the SWD in S1. B) LFP traces shows SWD-breaks in the indicated regions. Vertical dashed lines indicate the start and end of the SWD-break in V1. C) Duration of onset delays from S1. D) Rate of occurrence of SWD-breaks. E) Duration of SWD-breaks. F) Onset delay and SWD-breaks combine to form the parameter “SWC-Failure” (see Figure S1) and comparisons were made from V1 (black diamond) to every other channel. G, H) LFP traces recorded in the indicated cortical regions before (G) and after muscimol (H) infusion show an increased onset delay and SWD-breaks in M1 and V1, vertical black and blue dashed lines indicate SWD onset time in S1 and V1, respectively. I, J) Changes in SWC-Failure, calculated with respect to the pre-injection period (−60 to 0 min), in M1 (bright green circles/squares) and V1 (bright blue circles/squares) during muscimol injection in PO (I) and LP (J) are compared to those after aCSF injection (dull green and blue triangles). Vertical dashed line indicates the start of the injection. *p<0.5, **p<0.01, ***p<0.001.

### SWD were not present uniformly across the cortex

As SWDs are first visible in S1, ictal onset and offset times were defined using the S1 channel. During the ictal period, there were instances of brief interruptions of the SWD in all other recorded regions, hereafter referred to as SWD-breaks (measured as the number of SWD-breaks per ictal second) (Figure 1B). V1 was significantly more susceptible to SWD-breaks (0.05 ± 0.07 Hz) than M1 (0.002 ± 0.001 breaks/s, p<0.0001), LP (0.02 ± 0.007 breaks/s, p=0.0006) and PO (0.02 ± 0.006 breaks/s, p=0.001) (Figure 1D). V1 SWD-breaks had a duration of 0.85 ± 0.03s, which was not significantly different to that of any other channel (Figure 1E). To quantify the overall time that any one region did not express Spike-and-Wave-Complexes (SWCs) during a SWD, the onset delay was combined with the duration of all the SWD-breaks and any difference in offset times to generate the parameter ‘SWC-Failure (%)’ (Figure S1). Again, V1 had significantly higher SWC-Failure compared to the other cortical and thalamic regions (13.1 ± 2.3%, vs M1: 3.7 ± 0.7%, p=0.0001; vs LP: 6.1 ± 1.7%, p=0.0026; vs PO: 5.4 ± 1.1%, p = 0.001) (Figure 1F).

### HO nuclei facilitate cortical synchrony during SWD initiation and throughout

To understand the neural basis of SWC-Failure, and if HO nuclei are involved, we pharmacologically inactivated the LP (n=8) or PO (n=9) by infusing the GABA_A_ receptor agonist muscimol in either one of these HO nuclei (Figure S2). Local application of muscimol in either nuclei did not affect the overall number or duration of SWDs (Figure S3) but SWC-Failure in V1 was significant increased (range 17 - 27%) (Figure 1G-J). This was sustained up to 90 minutes after injection in both HO nuclei (aCSF: -3.6 ±1.8%, LP: 11.98 ± 4.1%, PO: 13.7 ± 3.7%, both p <0.0001). Infusion of muscimol into the PO also significantly affected M1 SWC-Failure for the first 60 minutes (30-60 mins: aCSF: -0.8 ±0.6%, PO: 11.1 ± 3.0%, p=0.0032). The change in SWC-Failure was due to significant increases in onset delay and number of SWD-breaks, and a non-significant increase in SWD-break duration was also observed after PO inhibition (Figure S4).

Fluctuations in V1 SWD power were temporally correlated with that of LP and PO, as quantified by the cross-covariance of ictal amplitude envelopes in all channel pairs (Figure 2A, B). The covariance coefficient for V1-LP (0.5 ±0.03) and V1-PO (0.6 ±0.03) was significantly higher than most of the other cortico-cortical and cortico-thalamic pairings (Figure 2B), demonstrating that variations in SWD power, and therefore neural synchrony, were correlated. V1 power changes just preceded that of both LP (65 ± 10 ms) and PO (41 ± 6 ms), however the LP or PO cross-correlation lag times were not significantly different to zero (p=0.19), or to each other (Figure 2C).

**Figure 2.**
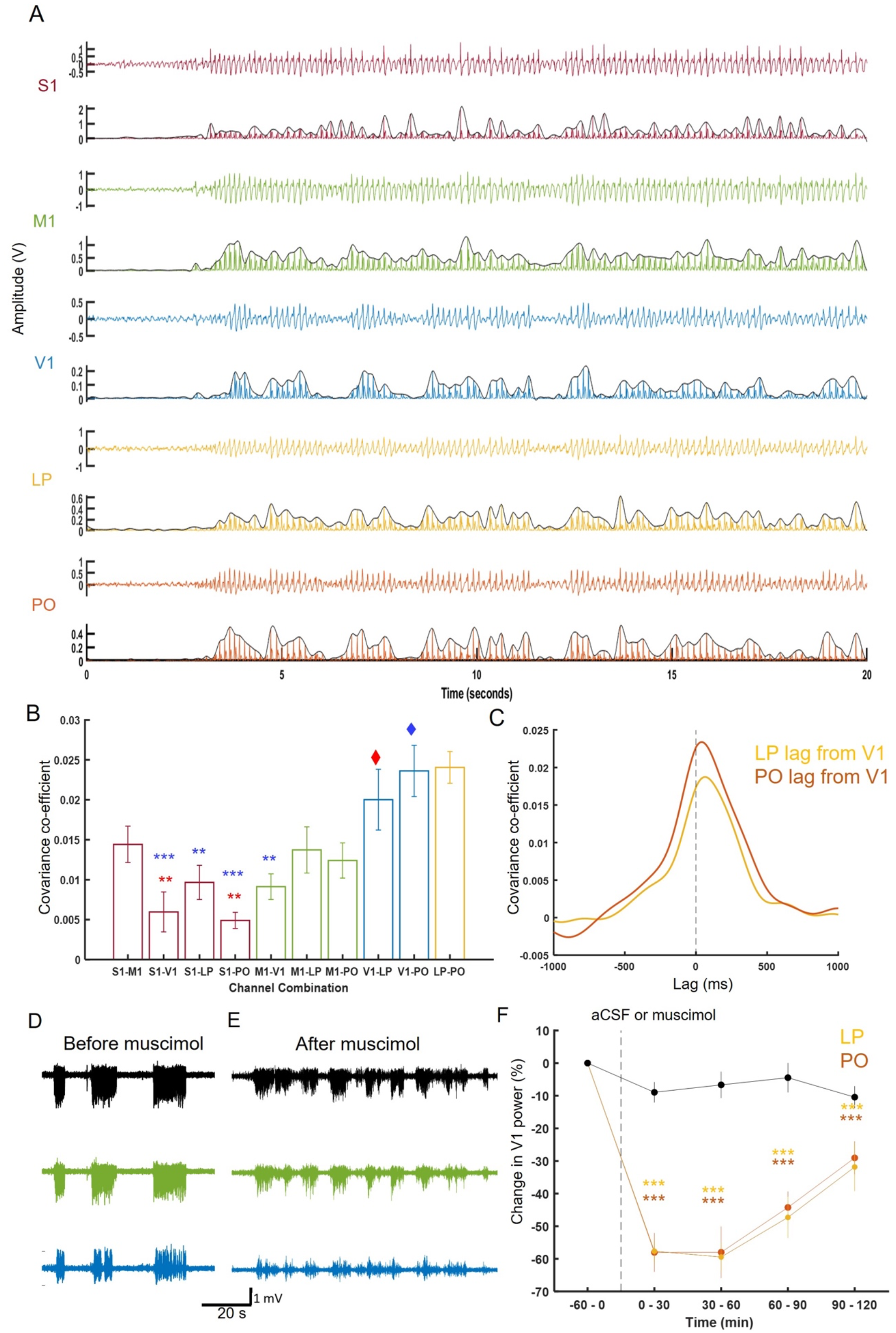
V1 SWD amplitude is temporally correlated with LP and PO and is reduced following inhibition of LP and PO. A) LFP (top trace in each pair) from cortical and thalamic regions and their squared signals with overlaid amplitude envelope (bottom trace in each pair) show that power of V1, LP and PO LFPs are more tightly correlated than other channel combinations. Cross-covariance was calculated on the amplitude envelopes and (B) coefficients were significantly higher in V1-LP (red diamond) or V1-PO (blue diamond) compared to most other channel combinations. C) There was a positive time-lag between LP and PO amplitude changes relative to V1. D, E) Example traces of S1, M1 and V1 before (D) and after muscimol (E) infusion showing a reduction in SWD amplitude. F) Changes in SWD power in V1 after muscimol infusion was significant larger when compared to aCSF injections. *p<0.5, **p<0.01, ***p<0.001.

Muscimol injection in LP and PO significantly affected the SWD power in V1 (animal and seizure, n = LP: 6 and 626; PO: 6 and 451), when compared to aCSF injection (animal and seizure, n = 6 and 668). The peak reduction occurred at 0-30 minutes post injection (aCSF: - 9.0 ±3.1%, vs. LP: 57.6 ±3.6%, p<0.0001, and PO: 58.0 ±6%, p<0.0001) and lasted until the end of the recording (Figure 2D-F). Muscimol injection into PO, but not LP, also affected the SWD power in S1 and M1, although this effect was more variable and not significant (Figure S5). As power reflects neural synchrony, these data show that V1 SWD synchrony is reduced by LP and PO inactivation, highlighting the involvement of HO thalamic nuclei in maintaining SWDs.

### HO thalamic nuclei contribute to successful SWD generalization

Successful generalization of SWDs is required for the behavioral manifestation of ASs. In our sample, SWDs were observed that occurred only in S1 (1.7 ± 0.7/hour), or in S1 and M1 (0.8 ± 0.3/hour), hereafter called “non-generalized events” (Figure 3A, B), whereas no similar events occurred in S1 and V1 only, or in S1 and LP/PO only. S1-only SWDs were 1.0 ± 0.06 s in duration. In S1+M1-only SWDs, S1 expressed a SWD for 1.3 ± 0.2 s and the M1 component started later and was therefore shorter (0.4 ± 0.1 s) (Figure 3C).

**Figure 3.**
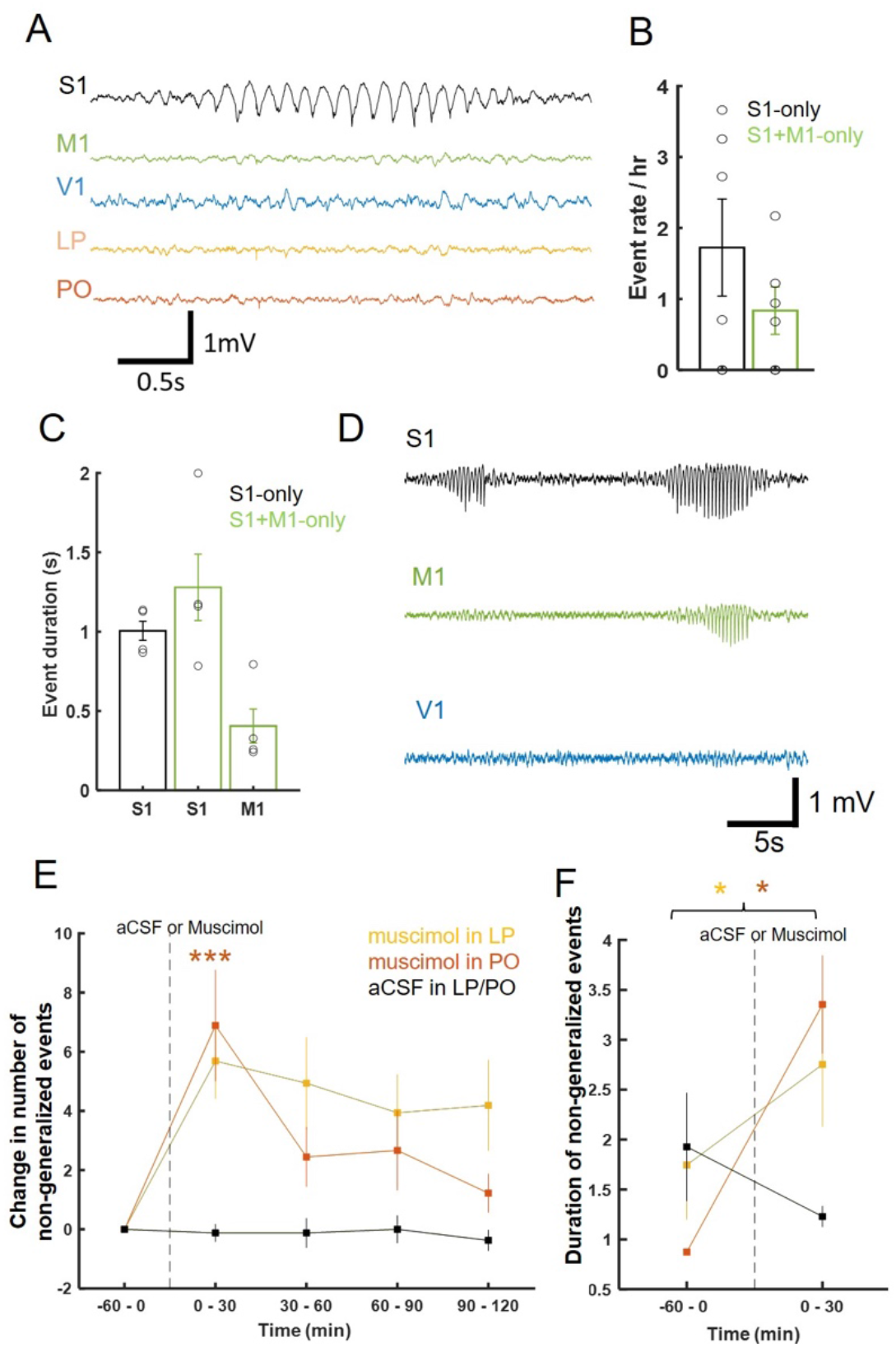
Features of non-generalized events (S1-only and S1+M1-only SWD) and effect of HO thalamic nuclei inhibition. A) Typical example of a non-generalized, S1-only SWD event in freely-moving GAERS. B, C) Quantification of rate (B) and duration (C) of non-generalized events. D) Typical example of a S1-only and a S1+M1-only SWD after muscimol infusion. E, F) Increased occurrence and duration (F) duration of both types of non-generalized events following muscimol infusion. Colour-code in (F) is the same as in (E). Dashed vertical line indicates the time of infusion of aCSF or muscimol infusion. *p=0.05, **p<0.01, ***p< 0.001.

There was a significant effect of muscimol on the occurrence of these non-generalized events since inhibition of PO and LP increased the occurrence of non-generalized events during the first 30 minutes post-injection (aCSF: -0.1 ±0.3 vs PO: 6.9 ±1.9, p=0.003, and LP: 5.7 ± 1.3, p=0.26), but only in PO was this change significant (Figure 3D, E). Moreover, the duration of S1+M1 non-generalized events was also affected by muscimol injections, with a significant increase observed after infusion when compared to their baseline durations (PO: 0.9 ± 0.03 to 3.4 ± 0.5 s, p=0.02; LP: 1.7 ± 0.5 to 2.7 ± 0.6 s, p=0.03).

### Optogenetic perturbation of PO activity suppresses SWD generalization

While muscimol infusions provided causal evidence that multiple HO nuclei contribute to SWD initiation and maintenance, this pharmacological intervention lacked temporal control. Therefore, optogenetics was used in 2 GAERS that had previously (3 weeks) been bilaterally infected in the PO (Figure S6) with a viral construct targeting local excitatory thalamocortical (TC) neurons (AAV5-CAMKII-ChR2-tdTom) to disrupt ongoing activity during SWDs.

Following single, brief (10 ms), bilateral 5.5mW pulses (n=40 trials) 5 seconds into a SWD, LFP rhythmicity was transiently reduced in all brain regions tested outside of the initiation site of S1 (Figure 4A) at a probability of 0.785 ± 0.035. To quantify the effect of optogenetic perturbation of PO activity, the correlation index of LFP (ratio of side peak and main peak in autocorrelations from 1s epochs before and following PO photostimulation, respectively) was calculated (Figure 4 B). Significant reductions were observed in M1 (pre: 0.86 ± 0.042, post: 0.69 ± 0.045, p=0.004) V1 (pre: 0.38 ± 0.01738, post: 0.13±0.072, p=0.00008), primary auditory cortex (A1) (pre: 0.53 ± 0.02027, post: 0.16 ± 0.0044, p=0.00008) and medial prefrontal cortex (mPFC) (pre: 0.91±0.036, post: 0.10±0.0032, p=0.00008). There was no change in rhythmicity of S1 LFP (pre: 0.91±0.036, post: 0.89 ± 0.044, p=0.13).

**Figure 4.**
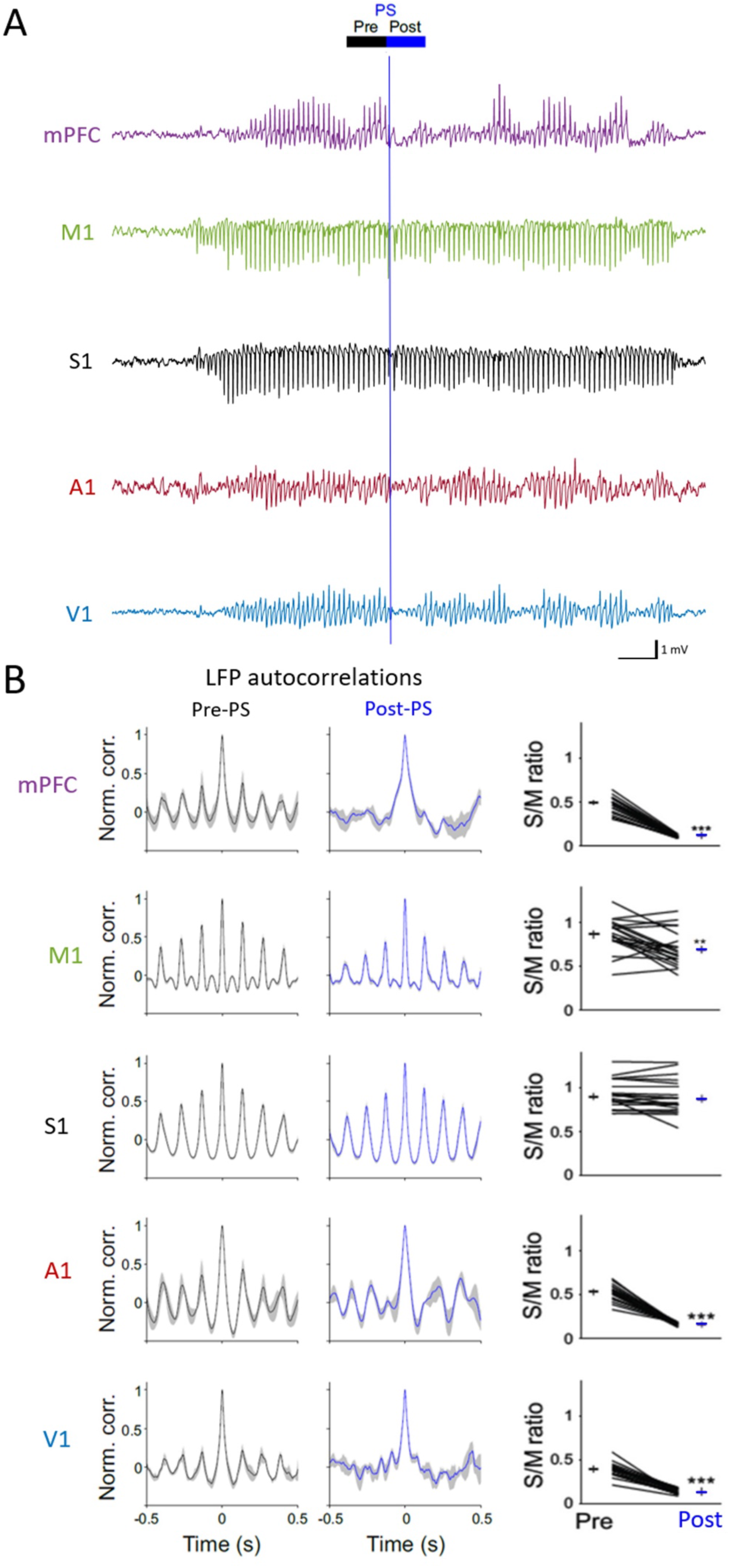
Optogenetic perturbation of PO activity results in SWD-break. A) LFP traces during a SWD showing the disruption of ongoing activity elicited by a single bilateral PO photo-stimulation (PS) 10 ms pulse in the indicated regions. B) Changes in rhythmic activity was quantified by calculating the Side-to-Middle-peak ratio (S/M ratio) of autocorrelations pre- and post-PS in the indicated brain areas (lower ratios indicate decrease of rhythmicity). **p<0.01, ***p<0.001.

### Two groups of PO neurons show different burst firing dynamics at SWD onset and during SWD-breaks

Since PO activity influenced SWD onset delays and SWD-breaks, we sought to understand the neural basis of these effects at the single unit level in this HO thalamic nucleus. A total of 94 PO units from 4 freely-moving GAERS were recorded with 4-shank silicon probes. As a population, the total firing rate began to decrease approximately 1.7 s before SWD start from 15.2 ± 1.1 Hz reaching a local minimum of 10.4 ± 0.6 Hz at 0.2 s before SWD onset (Figure 5A). Then, there was a rebound to 12.1 ± 0.8 Hz after onset, which was followed by a slower secondary decline to reach a stable ictal rate of approximately 9 Hz, within 3-5 s (Figure 5A). Interictal rates returned to pre-ictal values by 10s after SWD offset.

**Figure 5.**
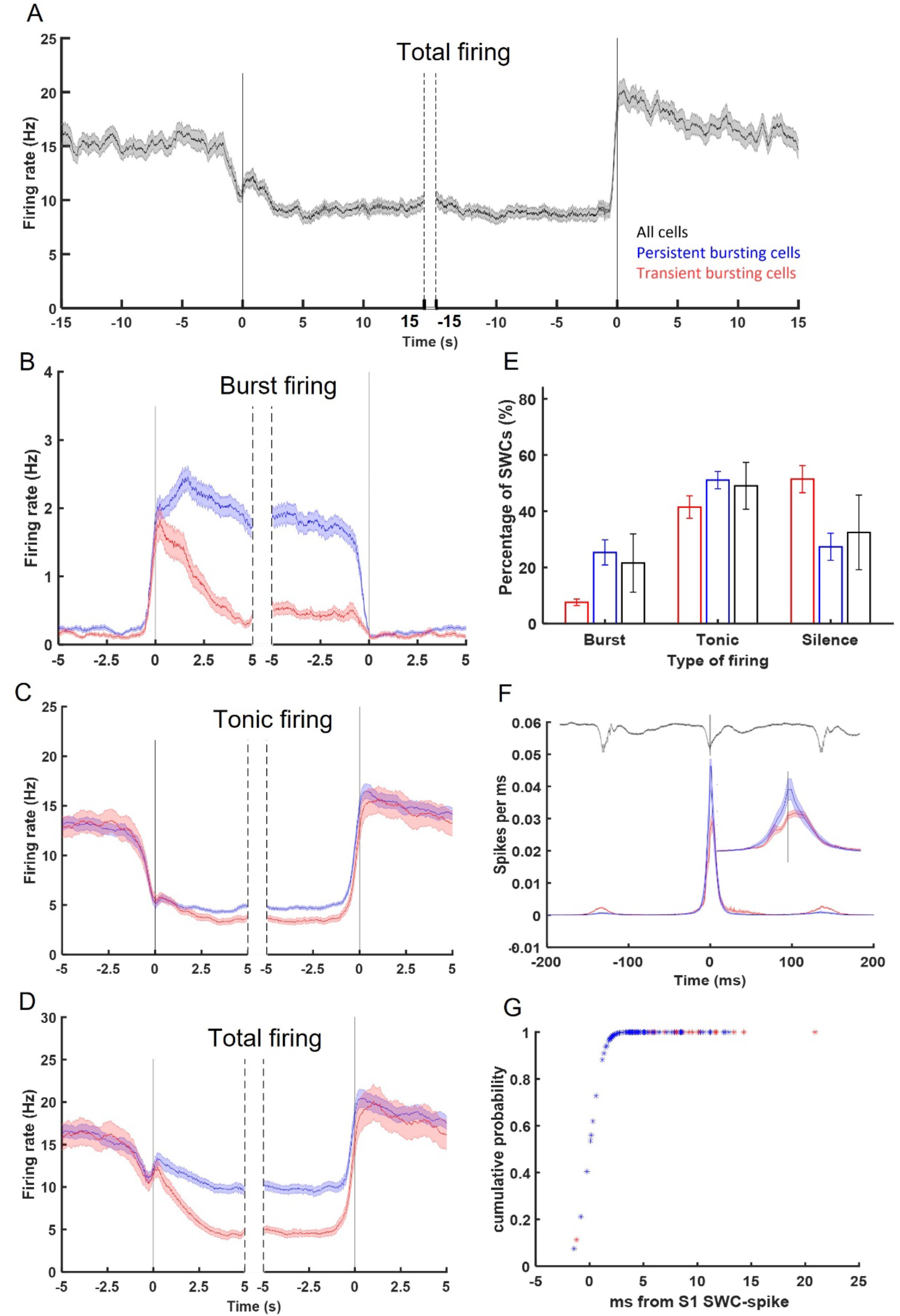
PO cells shift from tonic to burst firing at SWD onset and form two coalitions based on their ictal burst firing. A) Total firing of all cells relative to SWD onset and offset (black vertical lines). B) Two groups of PO cells, persistent (blue) and transient (red) bursting (PB and TB, respectively), were identified based on their burst firing dynamics. C, D) Tonic (C) and total (D) firing was similar between PB and TB cells. E) Percentage of SWCs showing tonic and burst firing or being silent F) Peri-SWC analysis shows a preference of PO neurons to fire after the SWC-spike in S1(top line is the corresponding LFP trace). G) cumulative probability distribution of peri-SWC firing. Color-code in E-G is the same as in A.

Unit firing was broken down into tonic and burst firing based on the classification in McCafferty et al., 2018 (see Methods). At SWD onset, the burst firing of one group of units increased to ∼2 ± 0.2 Hz and continued at this rate throughout the entire discharge: these neurons were named persistent bursting (PB) neurons (Figure 5B). The second group, named transient bursting (TB) neurons, had a similar increase in burst rate at SWD onset but then it reduced shortly after SWD onset from 1.7 ± 0.2 Hz to 0.5 ± 0.05 Hz which was maintained throughout the rest of the ictal period (Figure 5B). This dichotomy in PO ictal burst firing was quantified and cells were split in PB (n=64) or TB (n=30) groups using a gaussian mixture model (Figure S7A, B). Both cellular groups returned to pre-ictal burst firing rates (<0.25 Hz) by SWD offset. Although the firing dynamics differed, there were no significant differences in the action potential waveform (Figure S7C) or the intra-burst firing dynamics between PB and TP neurons (Figure S7D), and all units fired bursts with the classical “decelerando” pattern that is typical of excitatory thalamo-cortical neurons (Domich et al., 1986) (Figure S7D), though TB neurons had on average shorter bursts (Figure S7D).

The increase in ictal burst activity was balanced by a roughly simultaneous decline in tonic firing (Figure 5C). Both groups went from an inter-ictal tonic rate of ∼12.5 Hz to an ictal rate of less than 5 Hz. During the switch from tonic to burst firing there was a transient decrease in total firing from approximately 15 to 11 Hz (Figure 5D) which reached a local minimum of ∼10 Hz at ∼300 ms before SWD onset. PB cells had sustained firing at 9.8 ± 0.5 Hz whereas TB cells had a drop in total firing that reflected the burst firing pattern and reached an ictal total firing rate of 4.2 ± 0.5 Hz. Notably, despite the ictal switch from tonic to burst firing, tonic firing remained the dominant firing pattern during SWD onset and throughout and, additionally, there was a large incidence of silence, particular in the TB cellular group associated with many SWCs (Figure 5E).

Peri-SWC-spike histograms revealed that PB neurons preferentially fired action potentials from -1 to +3 ms with respect to the SWC-spike, whereas TB neurons fired from 0 - 5 ms (Figure 5F). A cumulative distribution function was also applied since it is less affected by variations in firing rate and this analysis showed both cellular groups were more likely to fire after the SWC-spike with only a 0.2 probability of firing before the SWC-spike (Figure 5G). The tonic to burst firing switch of PO neurons was less prominent at the onset of non-generalized events when compared to successful SWDs (Figure 6). Burst firing rates were lower at the onset of a non-generalized event (0.3 ± 0.08 Hz), compared to SWD onset (PB: 1.3 ± 0.1 Hz, TB: 1.2 ± 0.2 Hz, p<0.001) than at non-generalized event (Figure 6A, D), and tonic rates were lower at SWD onset than at non-generalized event, although this was not significant (Figure 6B, E). These differences lasted for about 1 s after onset, i.e. the approximate duration of a non-generalized event, and the total firing rate was the same in both events (Figure 6C, F).

**Figure 6.**
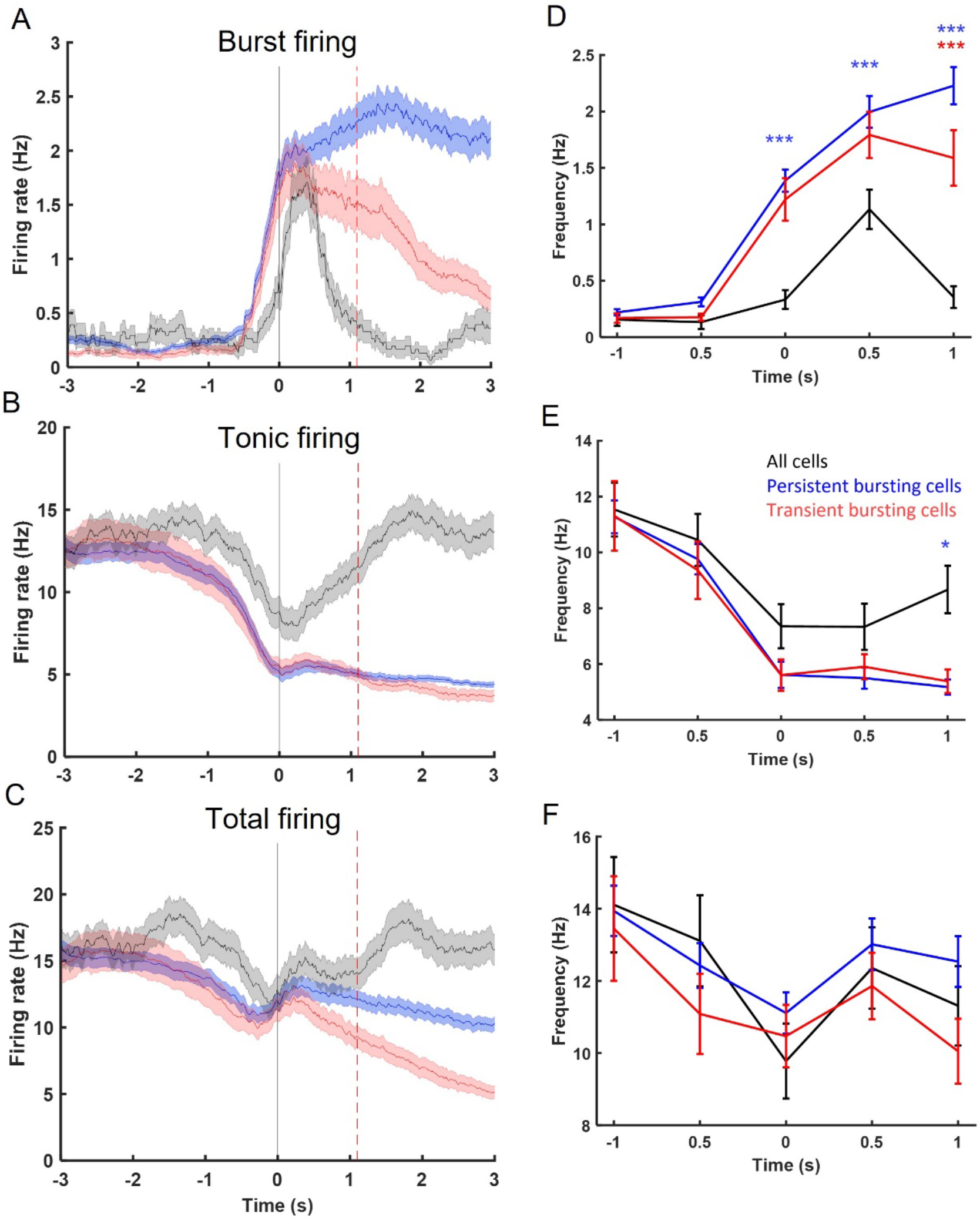
PO neurons fire less bursts at the start of non-generalized events. A-C) Burst (A), tonic (B) and total (C) firing for all cells and for PB and TB cells (vertical black and red line indicates the start and end, respectively of the averaged non-generalized events). D-F) Quantification of burst (E), tonic (E) and total (F) firing rates of PB and TB neurons at SWD-onset compared to population firing non-generalized events (black) for specific time points around onsets. Color-code in E applies to all other panels. *p=0.05, **p<0.01, ***p< 0.001.

The tonic to burst switch observed at SWD onset was transiently reversed during V1 SWD-breaks (Figure 7A-C). In PB neurons, burst firing declined from approximately 1 s prior to the onset of SWD-breaks from 1.58±0.6 Hz to 1.25± 0.5 Hz at break onset which was balanced by an increase in tonic firing. During this period, rhythmicity also changed, as observed by differences in autocorrelations obtained from spikes during the break compared to pre-break epochs of similar duration (Figure 7D, E). This was quantified by comparing the co-efficient at non-zero maximal peaks: the first maximum peak was highest at -2 to -1s and there was no difference between -1 to 0 s and the break period. The reduction in rhythmicity is exaggerated by the second side-peak of the autocorrelation with both pre-break epochs being higher than that of the V1 SWD-break. TB cells showed an increase in tonic and total firing at SWD-break onset and had low rhythmicity throughout: therefore, maximal peaks were not compared to V1 SWD-breaks.

**Figure 7.**
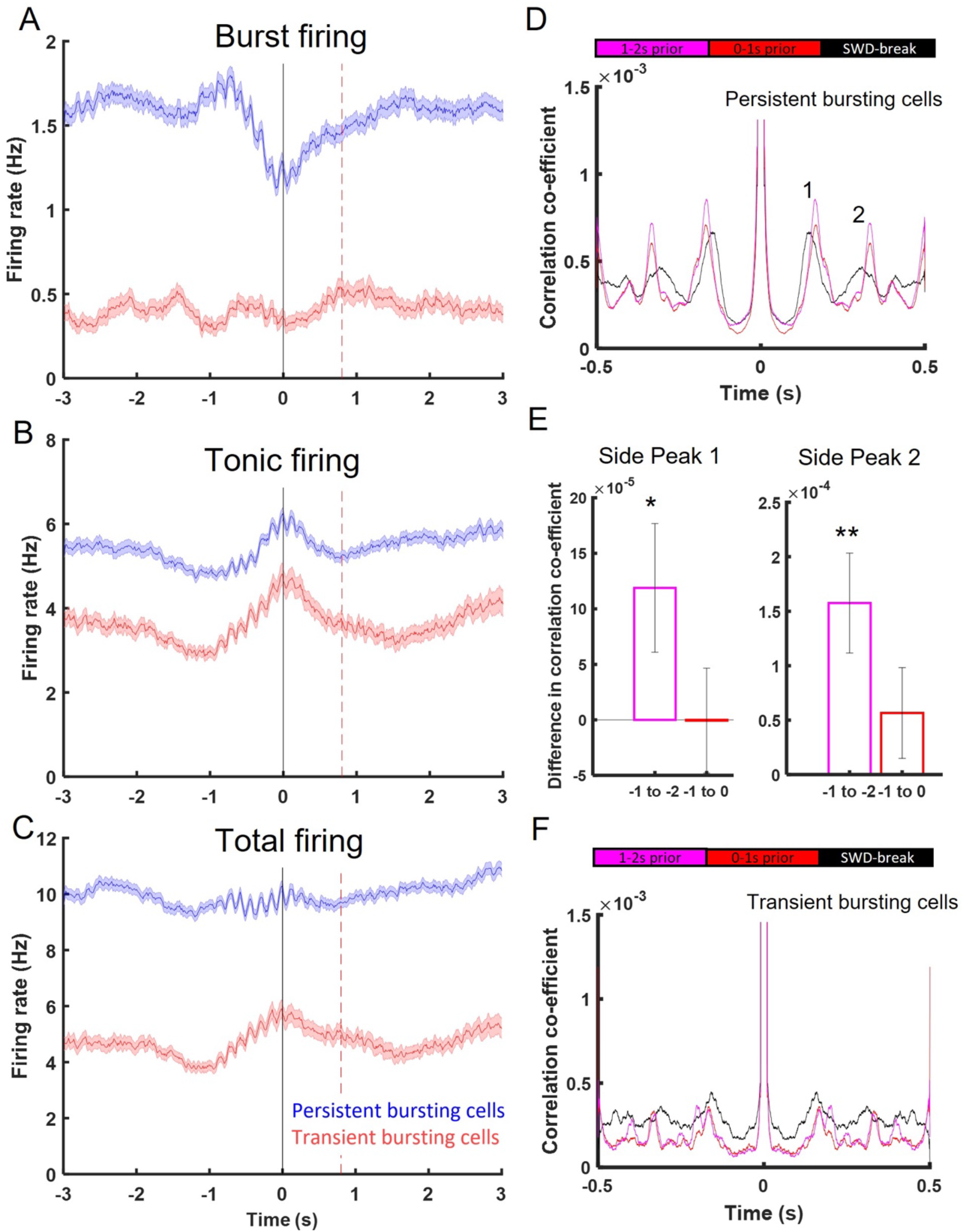
PO neurons change firing pattern, but not rate, at SWD-breaks. A-C) Burst firing of PB cells reduces at the onset of V1 SWD-break (A) while tonic (B) and total (C) firing changes less (vertical black and red line indicates the start and end, respectively of the averaged V1 SWD-break) (color-code in C also applies to A and B). D) The rhythmicity of PB neurons was lower during the V1 SWD-break. E) Quantification of the rhythmicity was performed by calculating the change in the maximum of side peaks 1 and 2 from the autocorrelation at SWD-break to the pre-break epochs in D. F) Rhythmicity of TB cells were relatively low in all epochs tested. *p<0.05, **p<0.01.

## DISCUSSION

This study provides causal evidence that HO nuclei are involved in the propagation and maintenance of generalized SWDs. Pharmacological manipulations via local muscimol infusions in LP and PO increase the SWD onset times and the occurrence of non-generalized events. Similarly, brief optogenetic perturbation of PO activity initiate SWD-breaks in various cortical regions. Interestingly, perturbations of LP and PO yield similar effects, despite the former not receiving direct input from the S1 initiation site. Furthermore, the neural basis of this influence was explored using single-unit recordings and we identified two PO neuronal sub-populations that differ in their ictal burst firing dynamics for the first time.

The contribution of PO to the generation of SWDs within the cortical initiation zone has already been indicated by observations of increased bidirectional coupling from a second before SWD onset and throughout (Lüttjohann and van Luijtelaar, 2015, 2012), potentially via direct excitatory reciprocal connections (Casas-Torremocha et al., 2019; Ohno et al., 2012; Veinante et al., 2000). Multiple studies show that PO stimulation results in excitation and sensitivity of cortical pyramidal neurons (Audette et al., 2018; Mease et al., 2016; Zhang and Bruno, 2019), which in GAERS, are intrinsically hyperexcitable (Polack et al., 2007). Additionally, there is evidence of an increased driver synapse efficacy from S1 to PO in GAERS (Çavdar et al., 2012) and thus, both S1 and PO could be more prone to generating recurrent excitation. However, our results challenge the necessity of S1-PO interactions for generating SWDs, as following PO silencing, the number of SWDs do not change significantly. This could be because PO has a facilitatory role rather than a critical one, resulting in more subtle changes such as the increased onset delays observed in our study. Alternatively, the number of PO neurons effected in this study may not have been sufficient to completely stop SWD generation, instead, the duration of SWD generation could have increased due to lower number of functional PO neurons. Notably, the intralaminar nuclei have recently been shown to provoke ASs under hypoxic conditions (Salvati et al., 2022) and thus even HO thalamic nuclei without recurrent connections with the initiation site can be important in SWD generation.

In addition to affecting SWD generation, there are multiple ways the HO thalamus can be facilitating cortical synchrony throughout the SWD. The spike component of SWCs represents synchronized cortical firing and when HO, but not FO, thalamic nuclei are inhibited, the spike component is reduced in regions outside the initiation site (Terlau and Luttjohann, 2020), a finding that strongly supports a wider synchronizing role of the HO thalamus. This is corroborated by the result of our study showing that LP and PO manipulations cause a strong reduction in cortical SWD power, another measure of neuronal synchrony. Furthermore, our study highlights that even HO nuclei that do not receive projections from S1, i.e. LP, are involved in SWD propagation and maintenance, further indicating that cortical synchrony is not occurring solely by cortico-cortical connections from the initiation site. These findings therefore argue that to accurately model the generalization of SWD, HO nuclei and their topographic cortical regions need to be included (see Martynas and Crunelli, 2023).

Topographic recurrent loops between cortical regions and their related HO nuclei is an emerging feature of cortico-thalamic networks (Guo et al., 2020). While PO-S1 excitation contributes to SWD generation, other thalamo-cortical modules do not have hyperexcitable pyramidal cells (Polack et al., 2007) and therefore these loops likely amplify rhythmic cortico-cortical excitation by entraining more neurons at every thalamocortical cycle until sufficient synchrony is obtained to result in an observable SWD. This process can explain the onset delays observed in our study. Although there is little overlap observed between these loops, such that S1 corticothalamic neurons project primarily to S1-projecting TC neurons in PO (Guo et al., 2020), there are PO neurons that do directly connect S1 to M1 and M2 via strong driver synapses (Mo and Sherman, 2019) as well as evidence of PO neurons projecting to both S1 and M1 (Ohno et al., 2012). This creates an additional opportunity of SWD propagation via trans-thalamic communication from the cortical initiation site to other cortical regions, particularly if the overlap zone is larger in GAERS, which is yet to be investigated.

Alternatively, HO nuclei can potentiate cortical input via projections to the apical tufts of layer V pyramidal cells located in layer I (Ledderose et al., 2021, Diamond ME., 1995). Activation of these dendrites is known to trigger back-propagating calcium spikes that can increase the activity of the pyramidal cells when coincident with feedforward cortico-cortical input (Larkum et al., 2001). Additionally, these superficial projections are not limited to specific cortical columns and thus provides a means of trans-thalamic communication and synchronization between thalamo-cortical modules. In these ways, each HO nuclei can be contributing locally to the integration of rhythmic cortical input during SWDs and the subsequent manifestation of an AS.

Our work provides the first insight of PO neuronal firing during SWDs in a freely moving GAERS. Sorokin et al. (2020) recorded from a mouse model of ASs and observed a ramp in PO firing preceding the onset, but this feature was variable and not observed here. However, overall firing rates were similar and alike Sorokin et al. (2020), a reduction in firing, followed by a sharp increase was observed just prior to SWD onset. Our work provides a major step forward to these results since we show that these dynamics represent an underlying shift from tonic to burst firing in the PO neuronal population.

We observed two sub-populations of PO neurons: TB neurons that fire bursts for the first second after SWD onset, the approximate duration of SWD generalization, and PB that fire bursts throughout the SWD. There was not a clear anatomical segregation between the groups as both cell types could be recorded during the same session and on the same shank. PO Connectivity and intrinsic cell dynamics are not homogeneous across PO, instead, there are distinct but overlapping input and output profiles (Casas-Torremocha et al., 2022, 2022; Guo et al., 2020) as well as highly varied burst patterns, that could explain the heterogeneity observed here (Desai and Varela, 2021). The complexity of thalamic neuronal dynamics at SWD onset extends to FO nuclei, where in VB neurons four groups of cells have been reported, including ‘onset peak’ and ‘sustained increase’ cells (McCafferty et al., 2021), similar to those reported here.

The tonic to burst shift observed in PO neurons is much weaker at the onset of non-generalized events and is reversed during V1 SWD-breaks, suggesting that the burst firing pattern is key for PO to facilitate SWD generalization. As the burst firing can generate larger EPSPs in its downstream targets (Hu and Agmon, 2016; Swadlow and Gusev, 2001), this may be a contributing factor to its associations with S1. These dynamics must be governed by an inhibitory input as hyperpolarization is necessary for burst firing. As S1 input is excitatory, this inhibition must occur via NRT or other extra-thalamic inhibitory afferents from the anterior pretectal nucleus or the zona incerta (Barthó et al., 2002; Bokor et al., 2005). Notably, the zona incerta has reciprocal connections with the initiation site and when inhibited, reduces SWD occurrence (Liang et al., 2011; Shaw et al., 2013). Although S1 has been shown to affect VB neurons via the NRT during SWD (McCaff), this cannot be generalised to the PO since NRT microcircuits between FO and HO differ in their location and properties (Martinez-Garcia et al., 2020).

In conclusion, our study provides causal evidence that HO nuclei, both those with and those without reciprocal connectivity with the cortical initiation site, are involved in SWD propagation. Therefore, the generalization of SWDs, and thus the manifestation of ASs, is not only determined by cortico-cortical connections, but also on more local thalamo-cortical modules and potentially trans-thalamic communication. These facilitatory mechanisms co-occur with a shift from tonic to burst firing of PO neurons and the complexity of thalamic neuronal activity during ASs is reflected in sub-populations of PO neurons. We now understand that SWD generalization and maintenance goes beyond the somatosensory loop and that there is a need for more detailed investigations into HO nuclei as a whole in the context of absence epilepsy.

## Acknowledgments

This work was supported by a Wellcome Trust PhD studentship to ZA (grant 204014/A/16/Z to VC), the Ester Floridia Neuroscience Research Foundation (grant 1502 to VC), the Hungarian Scientific Research Fund (Grants NN125601 and FK123831 to M.L.L.), the Hungarian Brain Research Program (grant KTIA_NAP_13-2-2014-0014), UNKP-20-5 New National Excellence Program of the Ministry for Innovation and Technology from the source of the National Research, Development and Innovation Fund to MLL. MLL is a grantee of the János Bolyai Fellowship.

