## Supplementary figures for "Higher-order thalamic nuclei facilitate the generalization and maintenance of spike-and-wave discharges of absence seizures"

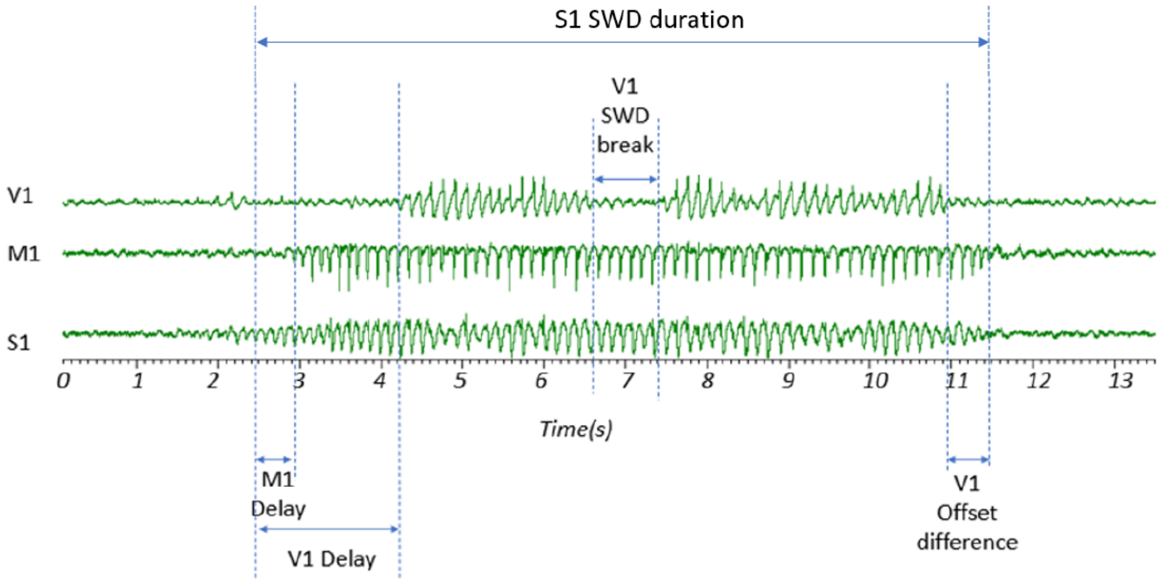


$$SWC \quad Failure = \left( \frac{V1\, delay + V1\, offset\, difference \; + \sum V1\, break}{S1\, SWD\, duration} \right) \text{x } 100$$

**Figure S1. Parameters that contribute to SWC-Failure.**

Raw traces illustrating the typical SWD waveform profile in S1, M1 and V1 which show an onset delay in V1 compared to S1, a SWD-break in V1 and a time difference at SWD offset in V1. These features combine to form SWC-Failure, as per the equation at the bottom of the traces. Note that, although not observed for this seizure, SWD delays, SWD-breaks and offset differences also occur in M1.

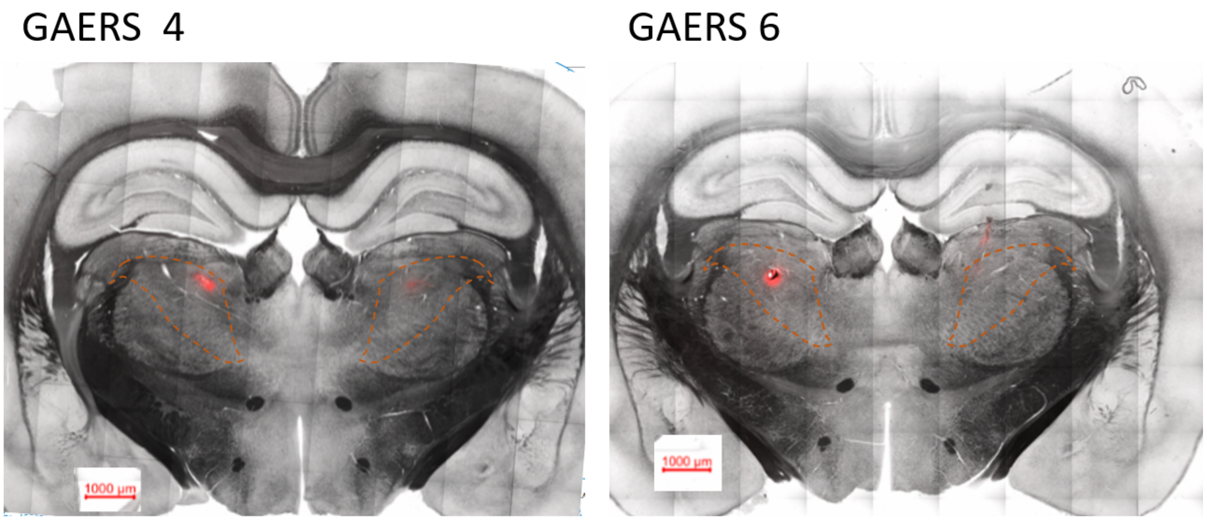


**Figure S2. Muscimol injection in LP and PO.**

Example histological slices demonstrating the spread of muscimol with a fluorophore conjugate in two different GAERS rats. Analysis of the signal via a confocal microscope at a fixed excitation power and pin-hole size enabled the signal to be measured and estimated to spread 200µM within HO thalamic nuclei as indicated by the contoured dashed red lines.

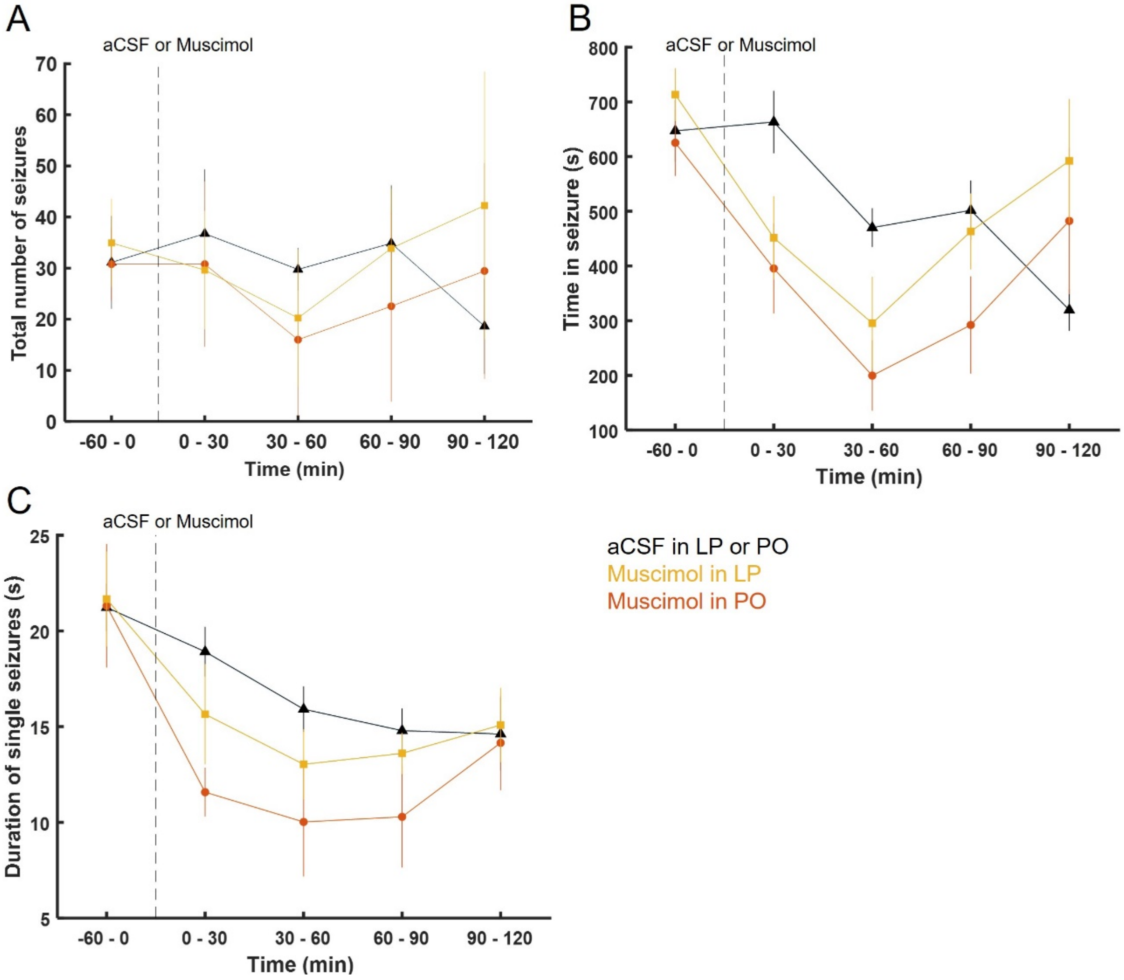


**Figure S3. Inhibition of HO thalamic nuclei by muscimol did not significantly impact basic seizure parameters.**

A) Total number of seizures, B) duration in seizures and C) duration of single seizures did not change significantly following muscimol injection in LP and PO compared to aCSF infusion at time 0 (vertical black dashed line) (color-code applies to A-C).

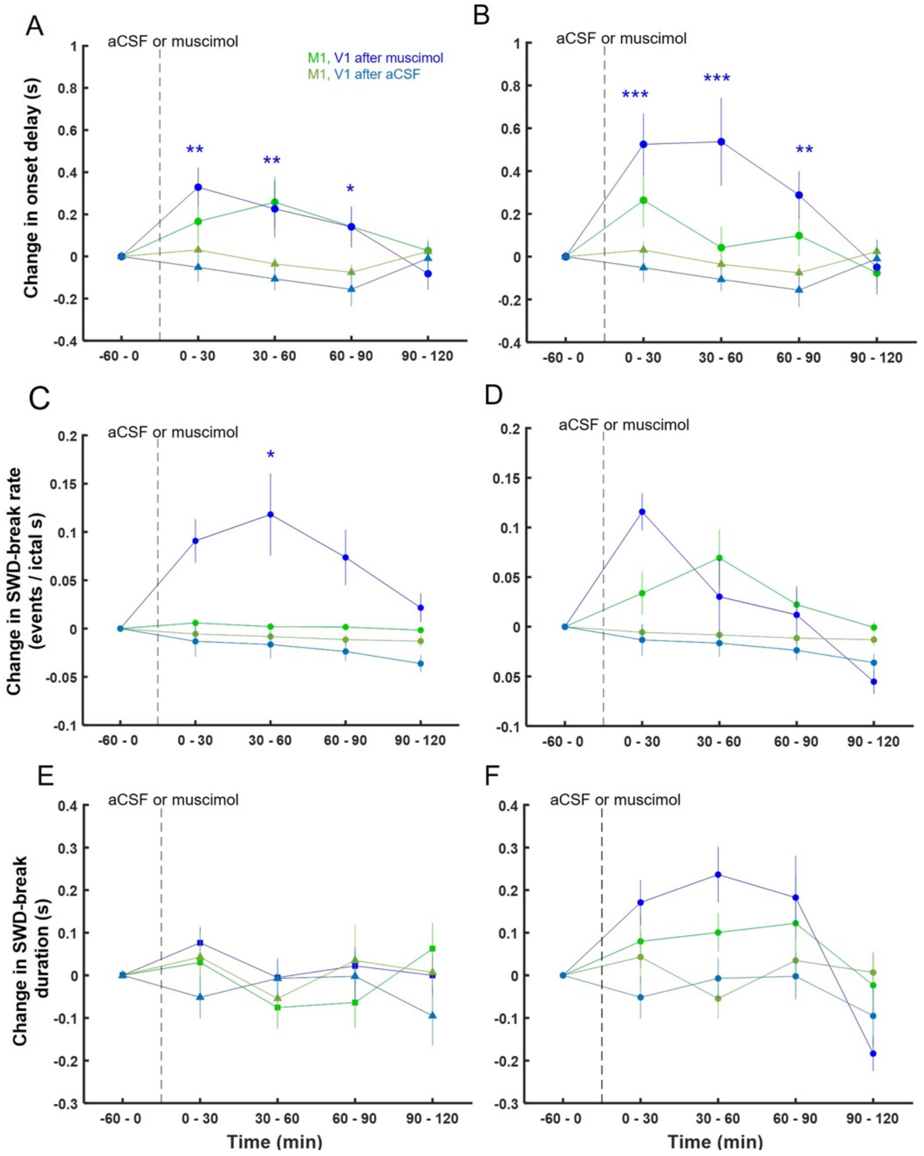


**Figure S4. Muscimol injection in LP and PO increased the onset delay and SWD-break rate in V1.**

A, B) Onset delay, C, D) SWD-break rate and E,F) SWD-break duration were differently affected in M1 (bright green lines) and V1 (bright blue lines) after muscimol injection in LP and PO compared to aCSF (dull green and blue triangles, respectively). Colour of * indicates the region significantly different to aCSF injections, either blue (V1) or green (M1). Black * indicates significant drug treatment F statistic. *$p<0.05$, **$p<0.01$, ***$p< 0.0001$.

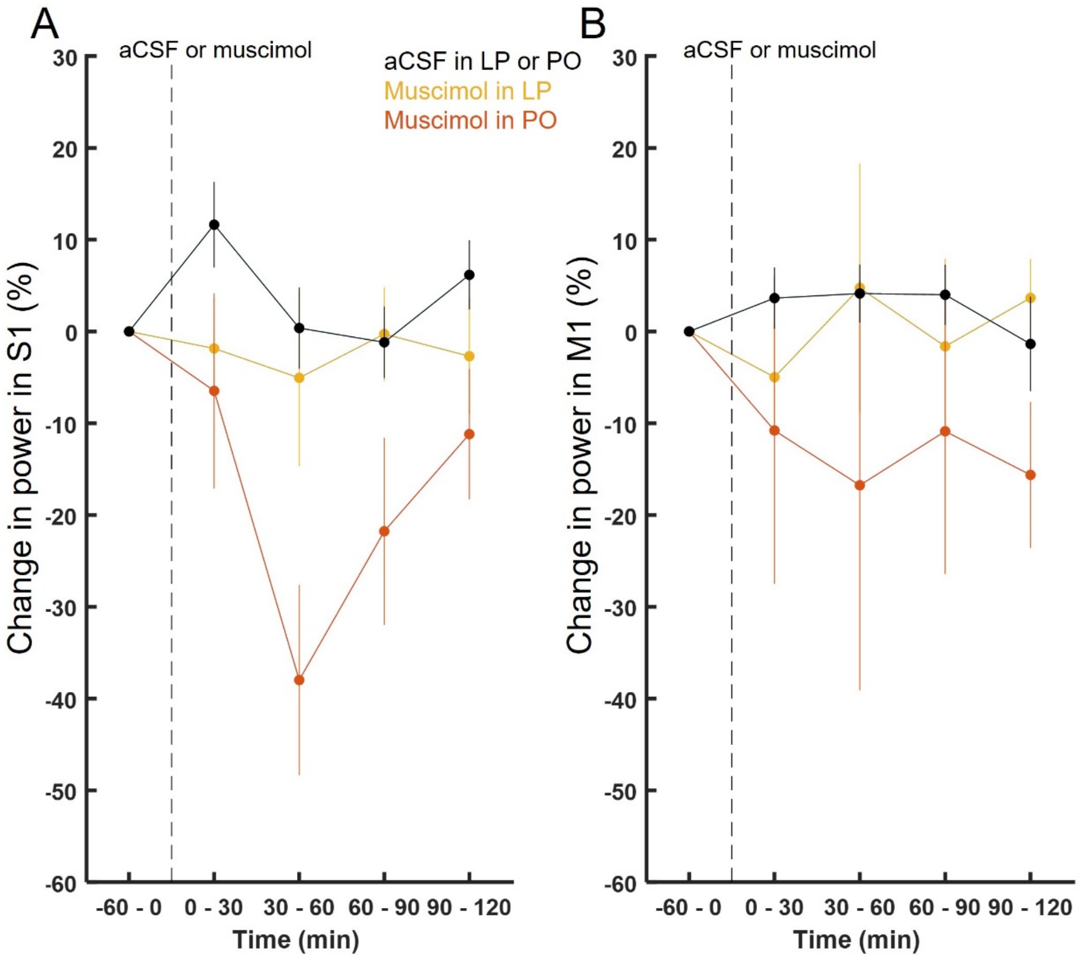


**Figure S5. Inhibition via muscimol infusion in LP and PO reduces SWD power in S1 and M1.**

A, B) Percentage changes in SWD power in S1 (A) and M1 (B) following muscimol injection in LP (yellow) and PO (orange) or aCSF injection in LP or PO (black).

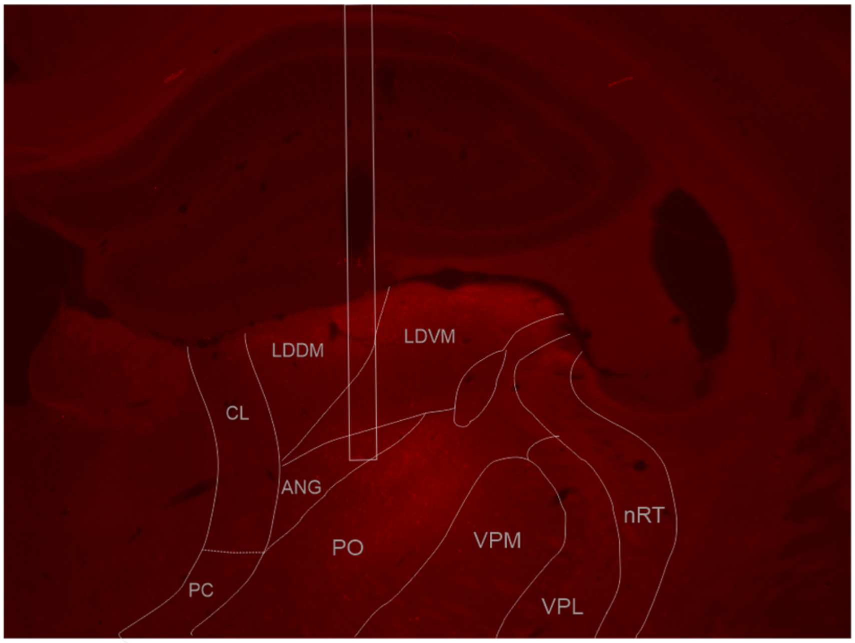


**Figure S6. Thalamic expression of ChR2-td tomato and location of the optic fiber.** Epifluorescent images (red channel: td tomato). Delineation of thalamic nuclei and optical fiber position are marked for clarity. Abbreviations: ANG: angular thalamic nucleus, LDDM: laterodorsal thalamic nucleus, dorsomedial part, LDVL: laterodorsal thalamic nucleus, ventrolateral part, CL: centrolateral thalamic nucleus, PC: paracentral thalamic nucleus, PO: posterior thalamic nucleus, VPM: ventral posteromedial thalamic nucleus, VPL: ventral posterolateral thalamic nucleus, nRT: nucleus reticularis thalami.

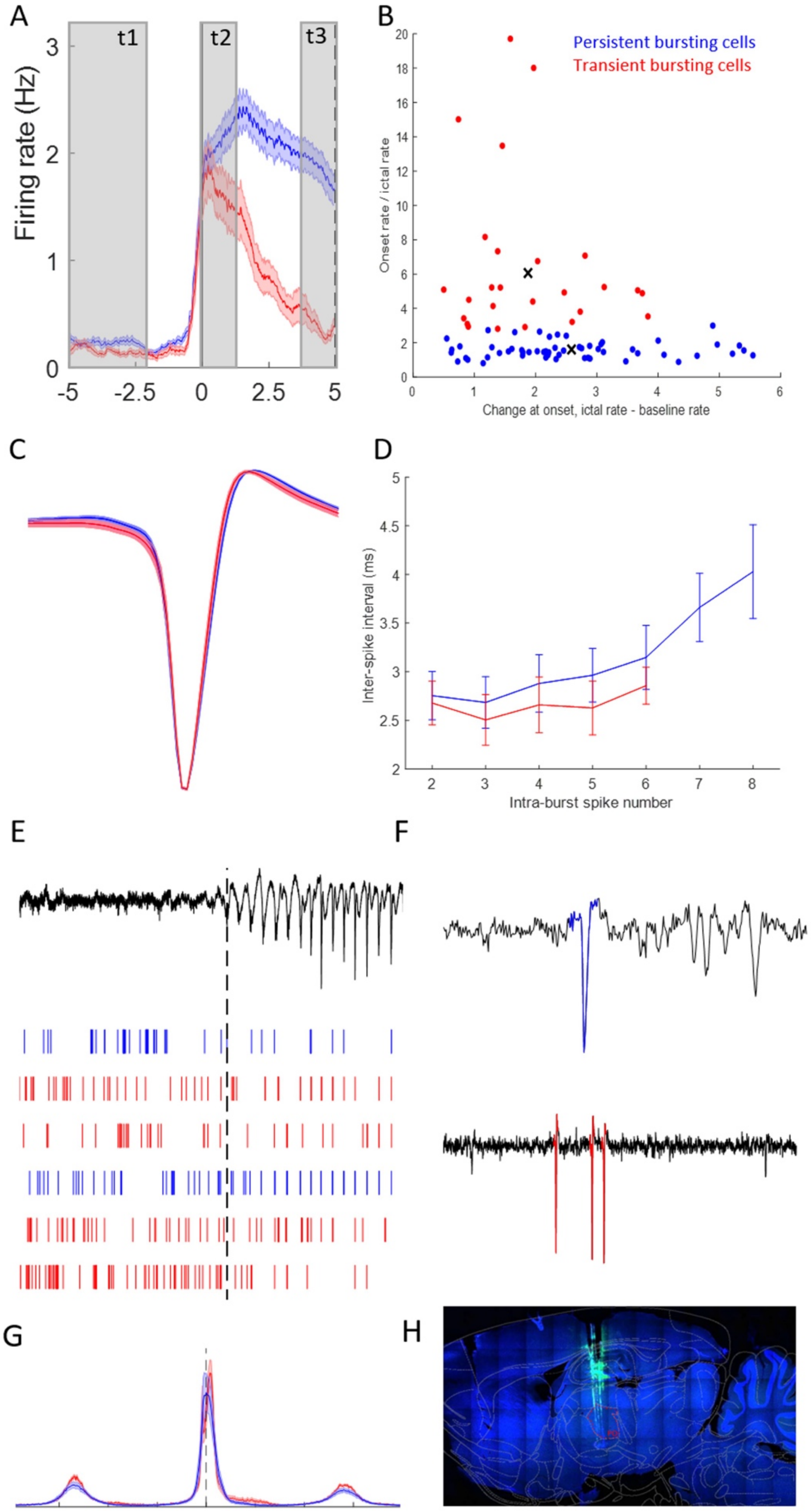
A
B
C
D
E
F
G
H
t1
t2
t3
Firing rate (Hz)
Persistent bursting cells
Transient bursting cells
Onset rate / ictal rate
Change at onset, ictal rate - baseline rate
Inter-spike interval (ms)
Intra-burst spike number

**Figure S7. Features of the two PO populations.**

A) Burst firing dynamics at SWD-onset (vertical line at SWD onset) of PB (blue) and TB (red) cells and epochs (highlighted in grey) used to calculate changes of firing rate at onset (t2 - t1) and onset rate/ictal rate (t2/t3). B) Those metrics were used to split up the total PO cell population using a Guassian mixed model into the two PO groups, color code applies to all panels. C, D) There were no differences in the extracellularly recorded action potential waveform (C) and burst firing pattern between the two groups. E) Example raster plots of cell firing at S1 SWD onset (dashed vertical line). F) Example raw traces with sorted spikes highlighted in blue (PB units, top) and red (TB units, bottom). G) There were no differences in firing-time relative to the ongoing SWC-spike between PB and TB PO cells. H) Example histology with silicone probe marked with DiI and depth marked with electrolytic lesions with traces from Watson and Paxinos atlas (1998) overlaid and PO highlighted in dashed red contour.
